# Locus-Specific DNA Methylation Editing in Mammalian Cells using a CRISPR-Based System

**DOI:** 10.1101/2021.10.12.463855

**Authors:** Jim Smith, Rakesh Banerjee, Reema Waly, Arthur Urbano, Gregory Gimenez, Robert Day, Michael R. Eccles, Robert J. Weeks, Aniruddha Chatterjee

## Abstract

DNA methylation is a key epigenetic modification implicated in the pathogenesis of numerous human diseases, including cancer development and metastasis. Gene promoter methylation changes are widely associated with transcriptional deregulation and disease progression. The advent of CRISPR-based technologies has provided a powerful toolkit for locus-specific manipulation of the epigenome. Here, we describe a comprehensive global workflow for the design and application of a dCas9-SunTag-based tool for editing a DNA methylation locus in human melanoma cells, alongside protocols for downstream techniques used to evaluate subsequent methylation and gene expression changes in methylation-edited cells. Using transient system delivery, we demonstrate both highly efficacious methylation and demethylation of the EBF3 promoter, a putative epigenetic driver of melanoma metastasis, achieving up to 304.00% gain of methylation and 99.99% relative demethylation, respectively. Further, we employ a novel, targeted screening approach to confirm minimal off-target activity and high on-target specificity of our editing sys-tem within our target locus.

## 1. Introduction

DNA methylation (5-methylcytosine; 5mC) is a stable, and perhaps the most widely studied, epigenetic modification involved in the regulation of gene transcription [1, 2]. Dysregulation of DNA methylation is implicated in the pathogenesis of numerous diseases. Aberrant DNA methylation in promoter regions of tumor-suppressor genes and global loss of DNA methylation has been strongly associated with the development and progression of many different tumors [3–5]. Classically, promoter DNA methylation is associated with transcriptional silencing [6]. However, several instances of promoter hypermethylation-induced gene activation have now been recorded [7–12]. We identified the *EBF3* gene as a putative epigenetic driver of melanoma metastasis [13] and in several other solid cancers [14], which shows paradoxical activation of transcription from highly methylated promoter. Understanding the precise mechanism of gene regulation via DNA methylation has great potential for advancing our understanding of disease pathophysiology and in identifying new targets for novel treatments [15]. Until now, it has been very difficult to establish true causality between DNA methylation changes and subsequent alterations in gene expression. However, with the advent of cutting-edge editing tools such as CRISPR, it is now possible to investigate the sequelae of aberrant methylation in diseases such as cancer in a causal manner [16, 17].

We have streamlined a method using the clustered regularly interspaced short palindromic repeats (CRISPR)-dCas9 system to facilitate site-specific editing of DNA methylation in mammalian cells [18, 19]. Although we have used this system to methylate and demethylate the promoter region of *EBF3*, using the CRISPR toolkit, the system described can be easily modified to target any known locus of interest within the genome, providing a highly selective method of epigenomic manipulation with minimal off-target impacts across the wider DNA methylome.

Our CRISPR-methylation system is based on an earlier system described by Huang et al (2017) [19], adapted for successful transient delivery into human melanoma cell lines and expanded to allow for targeted DNA demethylation alongside methylation. Immortalised cell lines are widely used as an experimental model for the fundamental investigation of tumour cell biology. DNA methylation status has been demonstrated to be well conserved between tumour tissue samples and derivative cell lines; therefore, cell lines provide an effective *in vitro* model for studying epigenetic alterations in cancer cells [4, 20, 21]. Our editing system comprises three main components: a dCas9-SunTag targeting protein; locus-specific guide RNA (gRNA; sgRNA) construct; and an effector construct for manipulating DNA methylation (Figure 1). The dCas9-SunTag construct is composed of a catalytically inactive *Streptococcus pyogenes* (*S. pyogenes*) Cas9 (dCas9) protein, fused to the SunTag (SUperNova TAGging) protein scaffold. dCas9 allows for RNA-programmable binding of our CRISPR-methylation editing system to a single target locus, without inducing cleavage of the underlying DNA sequence. Further, SunTag provides a repeating, epitope-based scaffold which is capable of binding multiple copies of our effector construct via short-chain variable fragment (scFv) domains [16]. dCas9-SunTag binding to a target genomic locus is directed by a unique gRNA construct. The *S. pyogenes* Cas9 module recognizes a 20 nt spacer sequence homologous to the target locus, which must immediately precede a 5’-NGG-3’ protospacer adjacent motif (PAM) [22]. Once targeted binding of dCas9-SunTag to our locus of interest has occurred, up to ten effector constructs bind to the SunTag scaffold via scFv binding domains. Here, effector constructs refer to proteins with the capacity to induce active methylation or demethylation of CpG dinucleotides, including the catalytic domains of the human DNMT3A methyltransferase or TET1 dioxygenase, respectively. The catalytic domain of the TET1 protein is preferred over the full length construct due to the difficulties with transfecting very large modules [23]. Collectively, these three constructs form our CRISPR-methylation editing system, with the capacity to induce active changes in DNA methylation at specific genomic loci. Here, broadly applicable protocols are detailed for gRNA design, and delivery of our CRISPR-methylation editing system into human melanoma cell lines.

**Figure 1.**
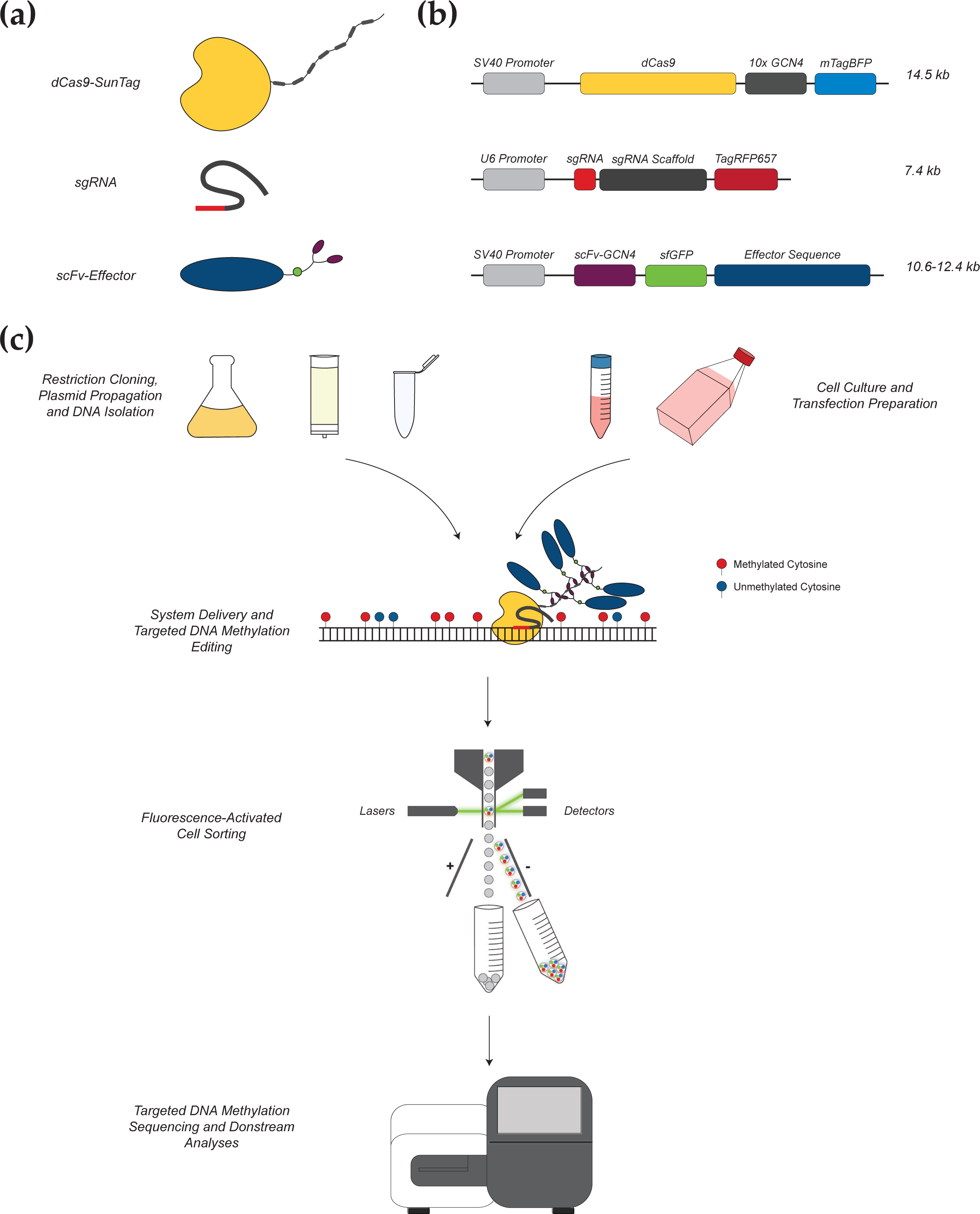
Overview of our CRISPR-based methylation editing approach. **(a)** Components of the CRISPR-methylation editing system. Shown are the three broad components of our editing system: a CRISPR-dCas9 construct for locus-specific targeting with an associated SunTag protein scaffold; a gRNA construct including a unique target sequence (red); and an effector protein construct (blue) with associated scFv domain (purple) for binding to the SunTag scaffold and tagged sfGFP fluorophore (green circle). **(b)** Structure and size of each plasmid is shown, corresponding to each of the respective constructs. **(c)** System components are cloned, propagated, and isolated as plasmid DNA for transfection into cultured cells, which provides transient delivery of each construct simultaneously to induce *in vitro* methylation or demethylation. A unique gRNA guides the CRISPR-dCas9-SunTag construct to bind at the target locus. Subsequently, multiple effector proteins bind to each respective GCN4 domain of the SunTag scaffold, wherein they can induce methylation change at surrounding CpG dinucleotides. Through modifying the effector enzyme, this system can be adapted for targeted DNA methylation or demethylation as desired. Post-transfection, cells positive for the expression of all components of the editing system are collected via fluorescence-activated cell sorting (FACS) and used for downstream analyses, including targeted DNA methylation sequencing. Adapted with permission from Urbano et al (2019) [16].

## 2. Materials and Methods

A full list of reagents and equipment for this protocol are detailed in Appendix A.

### 2.1 gRNA Design for CRISPR-Methylation Editing

The simple design of gRNA sequences for CRISPR experiments using the cloud based Benchling platform (http://benchling.com) has been previously described [24] and can be applied to any known target sequence. Benchling and other CRISPR gRNA design tools use an algorithm-based approach to generate potential gRNA sequences specific to a target locus with respect to PAM site requirements. In this protocol, we use the *S. pyogenes* Cas9 system, which will limit target gRNA sequences to those immediately preceding 5’-NGG-3’ [22]. Each algorithm then weights prospective gRNA sequences based on their projected on-target specificity and off-target activity. In CRISPR-methylation editing, minimising off-target activity is crucial to establishing causal roles for DNA methylation in pathways such as transcriptional regulation (see Section 2.5 for further discussion of off-target effects and an effective protocol for the targeted evaluation of off-target activity). When selecting gRNA sequences for methylation-editing, there are several additional factors that need to be considered. Firstly, as methylation-editing uses a dCas9-SunTag component, binding of dCas9 to the target locus will physically obstruct 20-30 bp directly overlying the gRNA target sequence [25]. The limited evidence currently available suggests that dCas9-SunTag-based systems are able to achieve efficient changes in DNA methylation up to around 1 kb from the PAM site [18]. Following design and gRNA selection, order oligonucleotides which represent both strands of the gRNA sequence; each oligonucleotide pair should have the following sequences:

> Forward oligonucleotide: 5’-CACCG(N)20-3’
>
> Reverse oligonucleotide: 3’-C(N’)20CAAA-5’

(N)20 denotes the unique user-designed guide sequence in the forward oligonucleotide and (N’)20 the reverse complement of the guide sequence in the reverse oligonucleotide. Each gRNA sequence requires, if not already present, the addition of a 5’ guanine residue (bold) which serves as a transcriptional initiation site for the U6 promoter in the final gRNA construct (a corresponding cytosine residue is therefore added to the reverse oligonucleotide sequence). A 5’-CACC sequence and 3’-CAAA sequence are added to the forward and reverse oligonucleotide, respectively, to generate complementary ‘overhanging’ sequences for ‘in-frame’ cloning into the BsmBI-digested pLKO5.sgRNA.EFS.tRFP657 vector. If possible, oligonucleotides should be ordered with pre-phosphorylated 5’ ends, removing the requirement for phosphorylation during gRNA construct preparation.

### 2.2 CRISPR-Methylation Plasmid Preparation

#### 2.2.1 Preparation of dCas9-SunTag and Effector Constructs

The dCas9-SunTag plasmid construct used here was obtained from Addgene (Catalogue #60903; pHRdSV40-dCas9-10xGCN4_v4-P2A-BFP). Construct preparation for the gRNA and effector proteins was based on the previously described protocols of Huang et al (2017) [13], using plasmids also available from Addgene. As per these vector construction protocols, pLKO5.sgRNA.EFS.tRFP657 (Catalogue #57824) was used as a vector for all unique gRNA constructs and pHRdSV40-scFv-GCN4-sfGFP-VP64-GB1-NLS (Catalogue #60904) was used as a vector for all effector proteins. All gRNA and effector constructs were prepared via restriction cloning, as per the cited protocols [19]. In brief, pHRdSV40-scFv-GCN4-sfGFP-VP64-GB1-NLS plasmids were restriction digested using RsrII and SpeI endonucleases overnight at 37 °C. Effector protein sequences were amplified from respective parent plasmids using primers with added RsrII and SpeI recognition sites and, subsequently, ligated into the digested vector. For gRNA construct preparation, pLKO5.sgRNA.EFS.tRFP657 plasmids were digested using BsmBI overnight at 55 °C then the ends dephosphorylated using rSAP (shrimp alkaline phosphatase). Respective forward and reverse gRNA oligonucleotides were then annealed and cloned into the digested vector (see Section 2.2.2).

For our work, all unique gRNA sequences were designed and selected as per the protocol detailed in the procedure section (see 2.1 gRNA Design for CRISPR-Methylation Editing). Two independent effector constructs were generated, containing sequences for the human DNMT3A protein and the catalytic domain of human TET1, respectively. The sequences for these effector constructs were derived from the following commercially available plasmids, respectively: Fuw-dCas9-DNMT3A (Catalogue #84476) and Fuw-dCas9-TET1CD (Catalogue #84475) [26]. It should be noted that if different plasmids to those stated are used for methylation-editing experiments, fluorophore selection within the plasmids is a key consideration to facilitate effective FACS (i.e., if using different plasmids, ensure any tagged fluorophores have sufficiently different emission spectra to allow for effective sorting of triple-positive transfected cells).

With respect to optional assessment of gRNA on-target editing specificity and potential off-target activity (see 2.6 gRNA Evaluation for On-Target Specificity and Off-Target Activity), our rapid screening protocol uses the active Cas9 construct pSpCas9(BB)-2A-GFP (Catalogue #48138) [27], also available from Addgene. This construct is propagated and isolated in the same manner as for the other plasmids used in this protocol. Simple screening of the isolated pSpCas9(BB)-2A-GFP plasmid DNA to confirm that the construct is of the correct size can be performed using EcoR1 restriction digest, which will generate two fragments of 8,505 bp and 783 bp, respectively.

#### 2.2.2 Preparation of Guide RNA (gRNA) Constructs

Begin gRNA construct preparation by performing restriction digestion of 25.0 µL (up to 10 µg) of the pLKO5.sgRNA.EFS.tRFP657 vector using 2.0 µL (20 units) of BsmBI restriction endonuclease overnight at 55 °C, in a total reaction volume of 105.0 µL, made up with appropriate enzyme buffer and water. Add 2.0 µL (2 units) of shrimp alkaline phosphatase to the digested product and incubate at 37 °C for 30 min, followed by 65 °C for 5 min to dephosphorylate the free ends of the digested vector. Purify the dephosphorylated vector using the DNA Clean and Concentrator-5 Kit or equivalent.

For annealing of each respective gRNA, combine 1.0 µL of sense oligonucleotide (100 µM), 1.0 µL of antisense oligonucleotide (100 µM), 1.0 µL of T4 DNA Ligase Buffer and 7.0 µL of water. Mix by pipetting and perform thermal cycling with the following protocol: 95 °C for 2:30 minutes; -1.0 °C per cycle for 10 seconds (repeat x 72); infinite hold at 22 °C. Dilute the annealed gRNA sample 1:500, then ligate into the digested pLKO5.sgRNA.EFS.tRFP657 vector at room temperature overnight using T4 DNA Ligase. Transform the ligated construct into competent *E. coli* cells for propagation on LB agar containing 100 µg/mL ampicillin, overnight at 37 °C. Confirmation of the inserted gRNA sequence can be performed via Sanger sequencing of single colonies using the U6 promoter primer (5’-TTTGCTGTACTTTCTATAGTG-3’), prior to bulk culture and transfection.

### 2.3 Transient Delivery of the CRISPR-Methylation Editing System

We recommend creating a transfection plan prior to each transfection experiment in order to establish reagent requirements and streamline the transfection process. In particular, detailed planning is important when performing complex transfections involving multiple constructs (i.e., when multiplexing gRNA molecules or using different effector constructs). This protocol describes a general method for the transient delivery of our CRISPR-methylation editing system into human melanoma cell lines via lipofection. It should be noted that other transfection methods may be better suited to different cell lines. Lipofection is performed using the Lipofectamine 3000 transfection system, with slight variation from the manufacturer’s protocol. Cells positive for successful plasmid delivery are subsequently sorted by FACS at 72-hours post-transfection.

#### 2.3.1 Cell Culture

For our optimized protocol, human melanoma cell lines WM115, CM150-Post and NZM40 were used. Cell line WM115 was obtained from America Type Culture Collection (Manassas, VA) (ATCC® CRL-1675TM). WM115 was cultured in Minimum Essential Media-Alpha (MEM-α) (Invitrogen) supplemented with 1% penicillin-streptomycin (Gibco, NY, USA) and 10% foetal calf serum (FCS). CM150-Post is a cell line established from patients entered into the Roche “BRIM II” phase II study of vemurafenib in patients who had previously failed treatment [28]. CM150-Post was cultured in Dulbecco’s Modified Eagle Medium (DMEM) (Invitrogen) supplemented with 10% FCS and 1% penicillin-streptomycin as previously described [3]. NZM40 was generously provided by Professor Baguley (University of Auckland). NZM40 was cultured in MEM-α supplemented with 5% FCS, 1% penicillin-streptomycin and 0.1% insulin-transferrin-selenium (Roche). All cell lines were cultured under standard conditions (5% CO2, 21% O2, 37°C, humidified atmosphere). Low passage and healthy conditions are essential to ensuring optimal transfection results. Cell lines were defrosted approximately one week prior to transfection, grown until >85% confluent in a 75 cm^2^ cell culture flask, then passaged to a 175 cm^2^ cell culture flask.

Melanoma cells were propagated in appropriate culture medium until >85% confluent. Appropriate culture medium and the length of time for cells to reach confluency will depend on the individual cell line. For best results, the following steps should be performed whilst the DNA-lipid complex(es) is/are incubating (see 2.3.3 Transfection). Trypsinise cells and transfer to an appropriate tube, then centrifuge for 5 min at 300 rcf. Remove culture medium and resuspend cells in an appropriate volume of culture medium for counting. Count cells and resuspend in culture medium to 5.0 x 10^5^/mL.

#### 2.3.2 DNA Preparation

Propagate each construct in *E. coli* cells overnight, with appropriate antibiotic selection. For our work, we use 200-500 mL of each respective culture in LB broth plus 100 µg/mL ampicillin, cultured overnight at 37°C, shaken at 200 rpm. Isolate plasmid DNA for each respective construct using an appropriate method. Isolating high-quality plasmid DNA with minimal bacterial endotoxin contamination is crucial for maximizing transfection efficiency and cell viability. Our preferred method for plasmid isolation is using the GenCatch™ Plasmid Plus DNA Maxiprep Kit. For convenience, these steps can be performed in advance and plasmid DNA can be stored at 4 °C until required for transfection. We recommend performing these steps up to several days in advance to minimise workload on the day of transfection. To confirm the identity of isolated plasmid DNA samples, Sanger sequencing or restriction digest may be used. For a simple restriction digest screen, it is possible to use the restriction enzymes which were used for cloning in reagent setup (e.g., RsrII/SpeI for effector protein construct(s)).

Prepare DNA combinations for transfection in sterile 1.5 mL microcentrifuge tubes by combining each respective construct in a 1:1:1 ratio, to the amount of 1500 ng total DNA. Each DNA combination must include dCas9-SunTag, plus gRNA(s), plus an effector protein construct. One or more unique gRNA molecules may be used (i.e., for targeting multiple genomic loci). If using multiple gRNAs, the total amount of gRNA DNA should still equal the DNA of each other construct (i.e., 500 ng total). For example, to perform targeted demethylation at locus X, we would produce a plasmid DNA combination containing 500 ng of our dCas9-SunTag, 500 ng of gRNA designed to target locus X, and 500 ng of our effector construct containing the TET1 catalytic domain. To target locus X and Y, we would instead use 250 ng of gRNA for targeting locus X, plus 250 ng of gRNA for targeting locus Y. For each run of transfections, include an empty tube which will be used for a negative control. Combinations can be prepared the day before transfection if desired. Mix combinations and spin briefly just prior to transfection.

#### 2.3.3 Transfection

Each step of the transfection process should be performed in a sterile cell culture hood. It should be noted that we use a reverse transfection method, which deviates from the standard procedure recommended by the manufacturer (i.e., resuspended cells are added to each well containing transfection reagents). First, add 5 µL P3000 reagent to 125 µL Opti-MEM Serum-Free Medium, per transfection reaction to be performed. Suspend each pre-prepared DNA combination in 130 µL of combined P3000 reagent and Opti-MEM Serum-Free Medium. Next, dilute 8 µL of Lipofectamine 3000 reagent in 125 µL of Opti-MEM Serum-Free Medium, per reaction, and add 133µL of combined Lipofectamine 3000 and Opti-MEM Serum-Free Medium to each suspended DNA combination. Incubate at room temperature for 15-20 min to allow for the formation of DNA-lipid complexes. For negative control samples, add all transfection reagents to an empty tube for transfection (i.e., as a negative control for FACS, cells will be exposed to all transfection reagents but no DNA, which provides a better control for FACS gating). Once each DNA-lipid combination has been incubated for 15-20 min, add each respective combination (263 µL) to an individual well of a 6-well cell culture plate. Next, add 1 mL of suspended cells in culture medium (5.0 x 10^5^) to each well. Add an additional 1-2 mL of culture medium to each well, bringing the total volume to 2-3 mL per well. Incubate under appropriate cell culture conditions.

#### 2.3.4 Cell Recovery and Maintenance

After the optimal exposure period, remove transfection reagents from the cells. The optimal exposure period to transfection reagents needs to be determined for each respective cell line and is defined as the length of exposure time to transfection reagents resulting in the highest percentage transfection efficiency without a substantial negative impact on cell viability (as assessed during FACS at 72 hours post-transfection). For our work, optimal exposure periods were 6 hours (WM115, CM150-Post) and 12 hours (NZM40), respectively. Wash with 1 x DPBS to remove any residual transfection reagent mix and replace with 2-3 mL of culture medium per well. Incubate under cell-specific optimal conditions until 72 hours post-transfection with changes of culture medium as required during this period.

### 2.4 FACS for CRISPR-Edited Cells

#### 2.4.1 Cell Preparation for FACS

At 72 hours post-transfection, begin cell preparation for FACS. Trypsinise cells, wash with 1 x DPBS and then stain with LIVE/DEAD Fixable Near-IR Dead Cell Stain Kit (Invitrogen) as per manufacturer’s instructions. After staining, centrifuge cells and remove supernatant, then resuspend in 250 µL of sterile autoMACS buffer. Also prepare and label a collection tube for each transfected sample (including negative control sample(s)) containing 250 µL of 1 x DPBS. Keep cells on ice until FACS is performed; for best results, perform FACS immediately once preparation is complete.

#### 2.4.2 FACS

Perform FACS using an appropriate system which allows for the simultaneous selection of cells based on positivity for multiple fluorophores. We recommend the use of the BD FACSAria Fusion (BD Biosciences) or similar. Our methylation-editing system requires the selection of triple-positive cells, exhibiting simultaneous positivity for mTagBFP, sfGFP, and TagRFP657. Begin the FACS run by running the negative control sample, which allows accurate gates to be set for identifying positive cells.

Collect negative control cells for later analysis. Once gates are set, sort and collect transfected samples into separate 15 mL Falcon tubes. Collect only live cells which are triple-positive (i.e., positive for mTagBFP, sfGFP, and TagRFP657 simultaneously; Supplementary Figure S1). Once sorted, centrifuge cells and remove supernatant, then store at -80°C until required for analysis.

### 2.5 Targeted DNA Methylation Analysis for Confirmation of CRISPR Methylation Editing

#### 2.5.1 DNA Preparation and Sequencing

Evaluation of CRISPR-edited cell lines and confirmation of successful DNA methylation editing can be performed using a variety of targeted DNA methylation analysis methods [18, 19, 29, 30]. We recommend a high-throughput, next-generation sequencing-based approach such as methylation-specific Illumina MiSeq sequencing [31]. Such methods pair bisulfite-specific sequence amplification with deep sequencing to offer high-resolution assessment of methylation status at a very low cost per read. Begin preparation for targeted DNA methylation analysis by extracting genomic DNA from CRISPR-edited cells and performing bisulfite conversion. For cell numbers >1.0 x 10^6^, we recommend the use of an appropriate extraction method for genomic DNA and subsequent bisulfite conversion using the EZ DNA Methylation-Gold Kit (Zymo Research). For samples with cell numbers <1.0 x 10^6^, we recommend the use of EZ DNA Methylation-Direct Kit (Zymo Research), which allows for cell lysis and bisulfite conversion without an intermediate DNA purification step. Comprehensive methodologies for next-generation [31] and other targeted DNA methylation analysis methods [32] have been published previously and, therefore, will not be detailed further in this publication. For bisulfite-specific primer design, we recommend using the MethPrimer online design tool [33]. It should be noted that the size of the genomic locus interrogated via targeted DNA methylation analysis will be limited due to bisulfite treatment (<500 bp). Illumina universal adaptor sequences should be added to the 5’ ends of the bisulfite-specific primers to facilitate sequencing on the Illumina MiSeq platform. Ordered oligonucleotide pairs should have the following sequences (where (N)22-25 is the bisulfite-specific primer sequence):

> Forward oligonucleotide: 5’-ACGACGCTCTTCCGATCT(N)22-25-3’
>
> Reverse oligonucleotide: 5’-CGTGTGCTCTTCCGATCT(N)22-25-3’

#### 2.5.2 Analysis of Targeted DNA Methylation Sequencing Data

Processing and analysis of methylation-specific sequencing data requires specific software packages in order to convert raw sequencing data into a useable output. We briefly outline here the software used in our analysis pipeline, specifically for Illumina MiSeq sequencing data of our methylation-edited samples. Analysis of targeted DNA methylation sequencing data has been described in detail by other groups [34, 35].

Briefly, the paired sequencing data should first be merged into full-length reads using PEAR (Paired-End reAd mergeR) software [36]. Merged reads then undergo quality assessment using FastQC (Babraham Bioinformatics) and subsequent adaptor trimming using Trim Galore! (Version 0.5.0, Babraham Bioinformatics). Processed reads can then be uploaded into the BiQ Analyzer HT software [37] in FASTA format, which aligns reads to a specified genomic reference sequence. BiQ Analyzer HT generates binary methylation data for each aligned read, at each CpG dinucleotide within the amplicon (Supplementary Figure S2).

### 2.6 gRNA Evaluation for On-Target Specificity and Off-Target Activity

Despite the continuing advancement of publicly available prediction algorithms and design software, definitive evaluation of on- and off-target activities for unique gRNA molecules can be a valuable adjunct to DNA methylation-editing experiments. Without the need for cost- and labour-intensive whole-genome approaches, we describe an optional protocol for rapid gRNA evaluation using an active Cas9-based assay.

#### 2.6.1 Selection of Predicted Off-Target Loci and Primer Design

First, select the top potential off-target sites for each respective gRNA. The most likely potential off-target loci are predicted via the *in silico* Benchling gRNA design process. We recommend selecting approximately twenty potential off-targets for evaluation. Subsequently, design a single set of primers for each off-target region using the NCBI Primer-BLAST online tool (https://www.ncbi.nlm.nih.gov/tools/primer-blast/index.cgi). Further, to establish the on-target specificity of a respective gRNA, design a set of primers that include the respective gRNA target sequence. Each primer pair should be designed to flank the respective on- or off-target locus plus >100 bp of sequence in either direction. Illumina universal adaptor sequences are added to the 5’ terminus of each primer to facilitate Illumina MiSeq sequencing. Ordered oligonucleotide pairs should have the following sequences (where (N)18-22 is the primer sequence):

> Forward oligonucleotide: 5’-ACGACGCTCTTCCGATCT(N)18-22-3’
>
> Reverse oligonucleotide: 5’-CGTGTGCTCTTCCGATCT(N)18-22-3’

#### 2.6.2 gRNA and CRISPR-Cas9 Transfection

Perform all steps of transfection and FACS as per sections 2.3 and 2.4; however, only transfect and sort a two-plasmid combination of gRNA (as prepared for methylation-editing) and active Cas9 (pSpCas9(BB)-2A-GFP). Perform genomic DNA extraction for each respective sample using an appropriate method. Do not perform bisulfite conversion here.

#### 2.6.3 Illumina MiSeq Sequencing

Again, preparation of Illumina MiSeq libraries for next-generation sequencing are well documented [31]. A multiplexed library should be generated using isolated DNA from transfected cells post-FACS and containing amplicons for each of the respective on- and off-target loci. The multiplexing strategy depends on the sequencing depth and the number of samples to be sequenced. Targeted amplicon sequencing libraries have low complexity, so PhiX genomic sequence is added to the sequencing pool to increase the diversity. The percentage of PhiX needed depends on the experimental design (i.e., the number of different regions to be sequenced). We obtained high coverage of 100x per sample together with minimum of 5% PhiX DNA, allowing reproducible methylation analysis. The sequence read length is dependent on amplicon size and where the target site is located within the amplicon. If possible, the amplicon should be centered on the targeted site. Read length should be less than the amplicon length to avoid read through the primer sequence. Paired end chemistry is the preferred option as it allows a more precise edition call, especially for insertion or deletion (INDEL) detection, due to the overlap of the two strands. The sequence overlap between paired end reads should be at least 10 nucleotides. Once sequencing is complete, data can be processed using the following command line-based analysis workflow.

#### 2.6.4 Analysis of Sequencing Data

First, sequencing reads with no mismatch compared to respective primers are demultiplexed using the grep command, then quality trimmed using fastq-mcf to include reads of ≥Q30 only. Overhanging sequences outside of the primers are then trimmed off, and Illumina universal adaptors removed using fastq-mcf. Paired-end reads are then merged (mergedPE) using FLASH (Fast Length Adjustment of SHort reads) [38], not allowing outies. MergedPE reads are then made unique (UniqSeq) and the number of occurrences is counted for each unique sequence. UniqSeq reads are aligned to the reference sequence using Needle (i.e., the Needleman-Wunsch global alignment algorithm).

Aligned UniqSeq reads are parsed, and single nucleotide polymorphism (SNP) and INDEL information retrieved for the respective in-target sites. The in-target site for each respective UniqSeq refers to the locus at which the sgRNA anneals to the reference sequence. SNP or INDEL (non-reference nucleotides) occurring at the in-target site is used as a surrogate measure of Cas9 activity and subsequent non-homologous end joining. True signal is distinguished from background noise via comparison against control samples (i.e., gRNA only and Cas9 only control samples). The below formula uses a conservative approach to estimate a background noise threshold using the maximum number of SNP or INDEL occurrences:

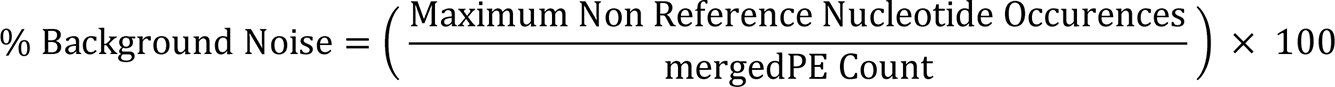

An ideal gRNA profile would, therefore, be supported by high occurrences of SNP and INDEL above background noise at on-target loci and with very low or no occurrences at off-target loci, respectively.

## 3. Results and Discussion

### 3.1 Locus-Specific gRNA Design for Targeted Methylation-Editing

Through careful gRNA design, we were able to achieve highly efficacious DNA methylation editing across three human melanoma cell lines, for a single target locus. Specifically, we targeted a 58 bp region of the *EBF3* gene promoter region (chr10:131,763,530-131,763,587; reference genome GRCh37/hg19), a gene that we have previously identified as a putative ‘epigenetic driver’ of melanoma metastasis [13]. This 58 bp locus, containing a total of nine CpG sites, was identified from genome-scale discovery analyses as a potential transcriptional control region and is located -993 bp from the *EBF3* transcription start site. We selected a total of six gRNA molecules from initial *in silico* gRNA design (Supplementary Table S1), comprising three distinct pairs, targeted 32-34 bp upstream (gRNA #1), corresponding to (gRNA #2), and 74-76 bp downstream (gRNA #3) of the target locus (Figure 2).

**Figure 2.**
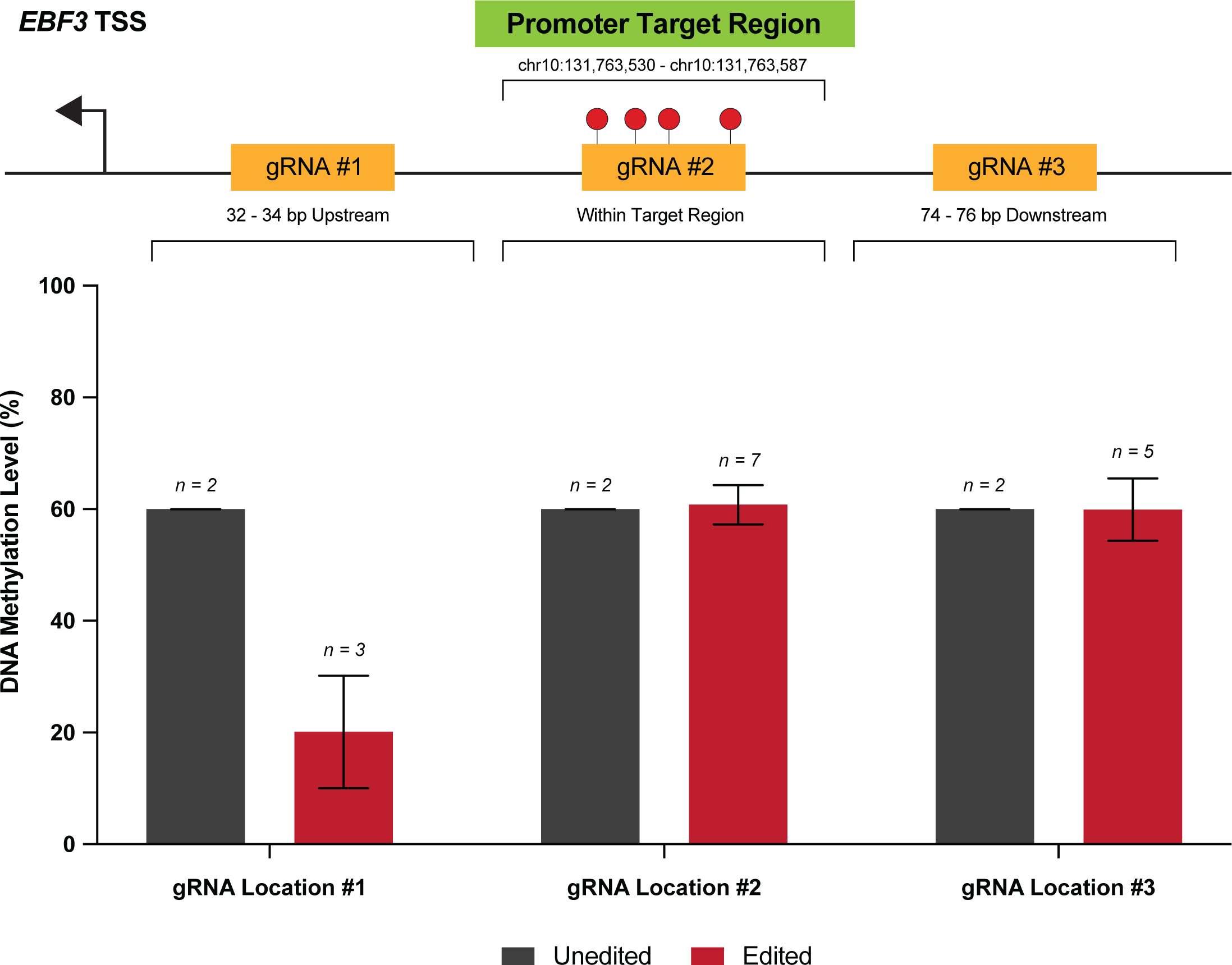
Variable efficacy of gRNAs designed to target the *EBF3* promoter. Shown are the three locations of the *EBF3* promoter, for which gRNAs were designed via *in silico* selection. For each location, two gRNA were designed, targeting the sense and antisense strands, respectively. Methylation levels within our target 58 bp locus are reported for cell line NZM40 in unedited versus edited samples, for each of the gRNA locations targeted. The position of each gRNA location with respect to the target locus is also shown.

**Table 1.** Pioneering studies in the development of CRISPR-based DNA methylation editing systems for mammalian cells.

| Group | System | Description | Features | Delivery | Stability |
| --- | --- | --- | --- | --- | --- |
| Vojta *et al* (2016) [29] | dCas9-Effector | Described the first dCas9 fusion with DNMT3A for targeted methylation editing in a ~35 bp wide region for the *BACH2* and *IL6ST* loci, with associated gene expression change | Simple design<br>Higher off-target activity<br>Moderate editing window (~150 bp) | Lipofection | Peak increase in methylation at day 6-7 post-transfection; persistent increases observed to at least 42 days |
| Xu *et al* (2016) [45] | dCas9, Tet1-MS2 | Designed a Effector-MS2 and dCas9 with modified gRNAs for effector multimerisation | Allows multimerisation of effector constructs<br>Lower editing efficacy<br>Moderate editing window (>100 but <300 bp; no editing within 100 bp of gRNA) | Polyethylenimine; Lipofection | mRNA expression change peaked 4 days post-transfection only |
| Morita *et al* (2016) [18] | dCas9-SunTag | Adapted the SunTag protein scaffold for multimerised DNA methylation editing, increasing the amino acid linker length to 22 nt to achieve demethylation efficacies of >90% both *in vitro* and *in vivo* | Allows multimerisation of effector constructs<br>Higher efficacy than dCas9-Effector, -MS2, -MQ1, -sMTase systems<br>Low off-target activity<br>Broader editing window (≤1 kb) | Lipofection | N/A |
| Lei *et al* (2017) [46] | dCas9-MQ1 | Fused dCas9 with an engineered prokaryotic CpG DNA methyltransferase “MQ1” derived from *Mollicutes spiroplasma* (*M.SssI*), strain MQ1 | Quicker onset of effective methylation change (24hrs)<br>Narrow editing window (~30-50 bp) | Lipofection | ≥ 3 weeks |
| Xiong *et al* (2017) [40] | dCas9-sMTase | Developed a targeted, “split methyltransferase” derived from *M.SssI* where the two split components bind at the target CpG site | Low off-target activity<br>Narrow editing window (8-25 bp) | Lipofection | N/A |
| Taghbalout *et al* (2019) [53] | *Casilio-ME* | Adapted the *Casilio* platform for methylation editing, providing co-delivery of TET1 and BER-associated proteins GADD45A or NEIL2 | Allows multimerisation of effector constructs<br>Comparable or higher efficacy than SunTag systems<br>Low off-target activity | Lipofection | N/A |
| Lu *et al* (2019) [47] | dCas9-R2 | Developed a dCas9-R2 module which specifically binds and inhibits the action of endogenous DNMT1 to prevent local DNA methylation at a target locus | Narrow editing window (<100 bp)<br>No requirement for exogenous effectors<br>Similar efficacy to dCas9, Tet1-MS2 | Lipofection | Peaked at 7 days post-transfection |
| Alexander *et al* (2019) [54] | Dual Cas9, Template | Excised a 1120 bp CGI from the *HPRT1* promoter using dual gRNA-guided Cas9 modules and replaced with either completely methylated or unmethylated fragments via NHEJ | Ensures complete de/methylation<br>NHEJ-mediated insertion has low efficiency (<1%)<br>Prone to repair-induced INDEL formation | Lipofection | N/A |
| Devesa-Guerra *et al* (2020) [39] | dCas9-ROS1 | Described a dCas9-Effector using the plant-specific DNA glycosylase ROS1, which directly excises 5mC | Higher gene reactivation efficacy than dCas9-Effector fusions alone; ineffective at fully methylated loci<br>Modest impact on measured methylation levels | Lipofection | N/A |
| Núñez *et al* (2021) [30] | CRISPROff | Developed a novel dCas9-Dnmt3A-3L-ZNF10 KRAB fusion (CRISPROff) to induce heritable, persistent gene silencing | High efficacy of methylation editing<br>Applicable to genome-wide screening<br>Stable editing effect despite transient delivery<br>Non-isolated effect of multiple enzymes (e.g., DNMTs plus KRAB) | Lipofection;<br>Nucleofection;<br>Lentiviral | Construct expression lost at 10 days; methylation change peaks at 9 days, stable for at least 30-50 days post-transfection |
| Smith *et al* (2021) (Current Manuscript) | dCas9-SunTag | Applied dCas9-SunTag with scFv-TET1CD and scFv-DNMT3A effectors to manipulate *EBF3* promoter methylation in melanoma | Achieved near-complete demethylation and high methylation<br>Comprehensive workflow for methylation-editing and downstream analyses | Lipofection | N/A |

Preliminary experiments were performed in cell line NZM40 to assess the efficacy of demethylation via dCas9-SunTag and scFv-TET1CD, using gRNAs targeted to each of these respective loci. Interestingly, whilst gRNA(s) targeted to gRNA #1 were able to induce successful demethylation at the target locus, no change in DNA methylation levels were observed for gRNA #2 or gRNA #3. Lack of response to targeting with gRNA #2 may be due to one of two reasons: (1) direct binding of the dCas9-gRNA complex is known to result in steric blockade of at least the underlying 20-30 bp of sequence, thereby precluding the action of bound effector proteins; (2) dense methylation at the target locus, alongside tightly-packed heterochromatin, may in itself mechanically interfere with dCas9 binding [25, 39]. gRNA location #3 may similarly be affected by inaccessibility to the DNA sequence, either at the point of dCas9 binding, or inaccessibility of bound effectors to target CpGs. Little is known with respect to the impacts of higher order chromatin organisation and topology on the action of DNA methylation editors; however, it is foreseeable that these factors have influenced the efficacy of our gRNAs for editing this particular locus. Indeed, achievable levels of editing efficiency and range of editing effect are variable between loci and editing systems, with ranges from as small as 8 bp to 1 kb from the PAM site currently reported [18, 40]. Hence, the design of multiple gRNAs is useful when targeting a specific locus. On the basis of these findings, only gRNA(s) targeted to location gRNA #1 were used for further methylation-editing experiments.

### 3.2 Efficient Locus-Specific Editing of the EBF3 Gene Promoter

Employing our aforementioned dCas9-SunTag system, in conjunction with scFv-DNMT3A and scFv-TET1CD effectors, respectively, we were able to induce both successful methylation and demethylation of the *EBF3* target locus across multiple cell lines (Figure 3). We chose a panel of three human melanoma cell lines with variable levels of baseline methylation at our target locus. Only one group has previously demonstrated epigenetic editing in melanoma cells, limited to modifying histone phosphorylation of the A375 cell line [41].

**Figure 3.**
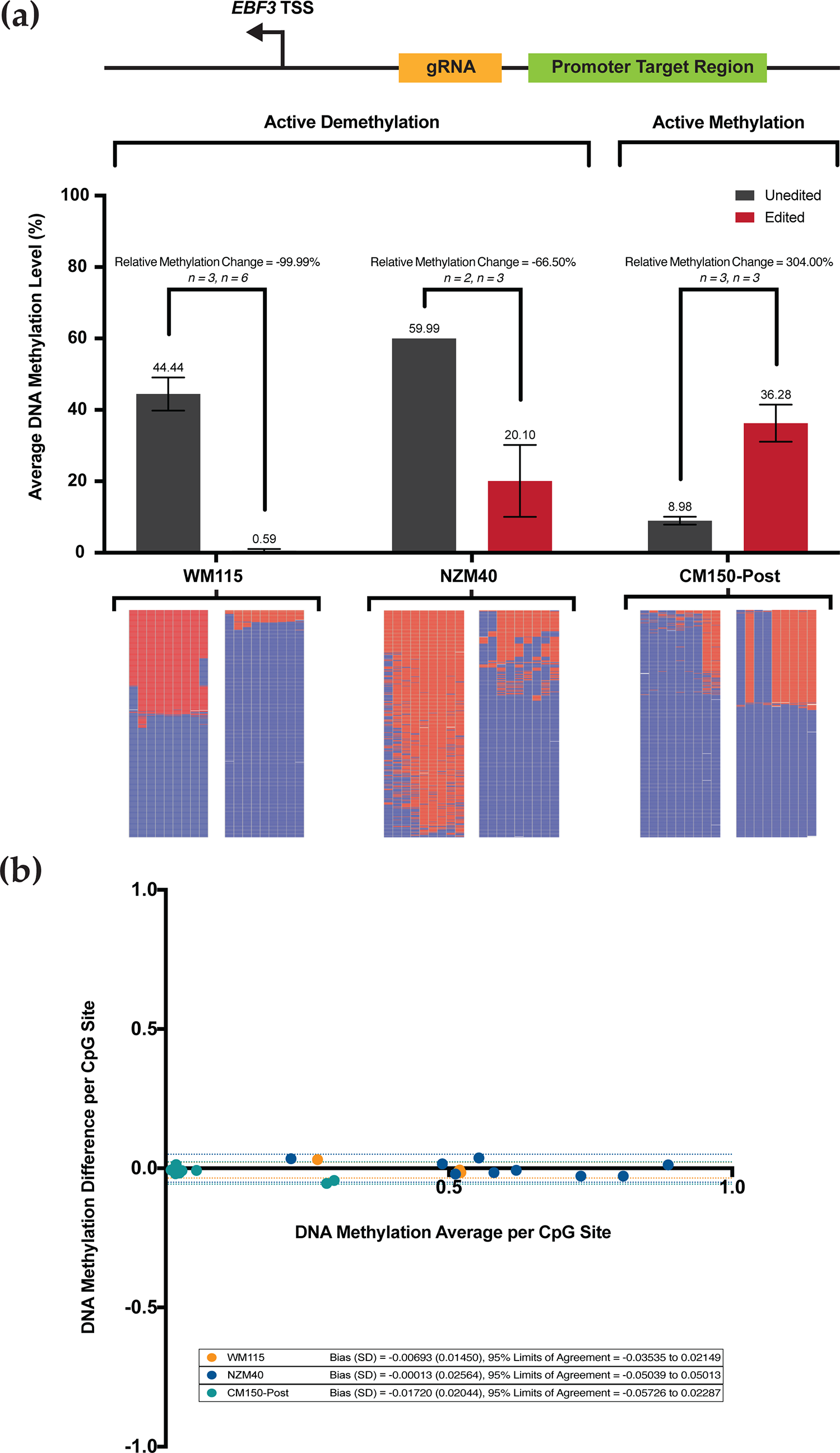
**(a)** Targeted DNA methylation and demethylation of the *EBF3* promoter region across three human melanoma cell lines. Average levels of DNA methylation (%) ± SEM (standard error of the mean; error bars) are shown for each of the cell lines WM115, NZM40 and CM150-Post, respectively, for both unedited and methylation-edited samples (absolute values per sample as shown). Note, no error bars are shown for NZM40 unedited sample as the SEM is too small to represent graphically. Active demethylation is demonstrated in WM115 and NZM40, whilst active methylation is demonstrated in CM150-Post. The relative level of DNA methylation change from unedited to edited is displayed as shown (%). Corresponding heatmap representations are shown for each sample, each of which was generated using 500 randomly selected sequencing reads to illustrate the methylation status of each CpG site within the target region; methylated (red), unmethylated (blue), or unaligned (white). **(b)** Bland-Altman analysis of replicate WM115, NZM40 and CM150-Post unedited samples per CpG. Plotted results of the Bland-Altman analysis between the corresponding per CpG methylation levels of successive replicate samples. Two technical replicates of unedited samples were analysed for each cell line as shown.

We performed targeted DNA demethylation in cell lines WM115 and NZM40, which displayed a relative methylation decrease of 99.99% and 66.50%, respectively (absolute methylation level change from 44.44% to 0.59% in WM115, and 59.99% to 20.10% in NZM40). Targeted DNA methylation was induced at this same locus in CM150-Post, resulting in a relative methylation increase of 304.00% (absolute methylation change from 8.98% to 36.28%). Relative methylation change between paired unedited and edited samples was calculated as:

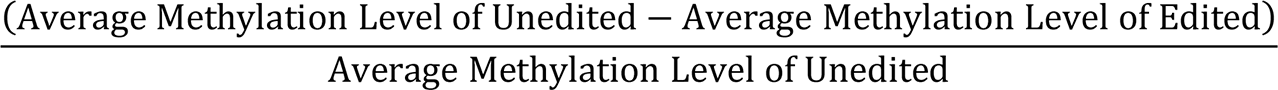

Biologically, these respective alterations in average DNA methylation level reflect a substantial change in the methylation status of the target locus between unedited and edited samples, which is consistent across all cell lines and technical replicates.

Interestingly, we observed more extensive changes in demethylation between WM115 and NZM40 cell lines, both relative and absolute. For cell line WM115, near-complete demethylation across the target locus was observed, whereas 20% of the methylation across this locus was preserved in NZM40. Previously, studies have demonstrated a difference in methylation-editing efficacy between cell lines due to disparities in transfection efficiency [42]. Using our approach, however, stringent FACS selection of only cells actively expressing our editing system, ensures that variations in transfection efficiency are adjusted for, providing directly comparable results between cell lines. Therefore, the reasons for this difference in our observed editing efficacy remain unclear, though an incubation time of 72 hours may have been insufficient to facilitate complete demethylation in NZM40. TET-mediated demethylation involves intermediate hydroxymethylation of cytosine residues, which is indistinguishable from 5mC via bisulfite conversion [43]. Hence, hydroxymethylated residues would be identified as 5mC during targeted sequencing. Slower-replicating cell lines would therefore undergo less complete 5mC loss via passive dilution in a given time frame and cells with lower expression of base excision repair-associated machinery may require additional time to achieve extensive methylation loss [44]. Each of these mechanisms may contribute to the differences observed in our cell lines, amongst other cell line-specific factors. With respect to gain of methylation in CM150-Post, the absolute mean methylation increase (27.3%) was modest in comparison to our demethylation experiments at the same locus. This increase did, however, represent a large relative increase (304.00%) in methylation, suggesting that it is likely to be significant from a biological standpoint. This result may reflect an innate resistance to gain of methylation at this locus for CM150-Post and it is plausible that active demethylation machinery may counteract the action of scFv-DNMT3A. Interestingly, Huang *et al* demonstrated a similar limit of editing efficacy with transient delivery of scFv-DNMT3A, though efficacy increased to around 80% over time when delivered via lentiviral transduction [19].

Overall, we report highly efficient and reproducible DNA methylation editing of a target *EBF3* promoter locus across a panel of human melanoma cell lines, using transient delivery of a dCas9-SunTag-based editing system.

### 3.3 Highly Reproducible DNA Methylation Analysis using Targeted Sequencing

Bland-Altman analysis (Figure 3b) was performed to asses*s* the reproducibility of successive sequencing replicates, as performed on unedited samples for each cell line. Note that for this analysis, DNA methylation level is reported as a proportion between 0.0 to 1.0, rather than as a percentage. This method compares the DNA methylation difference (y-axis) versus average (x-axis) between two successive unedited samples for each respective CpG site (i.e., CpG 1-9 across the target region), for each respective cell line. The bias values, which represent the average of plotted differences for each respective cell line, are very low, at -0.00693 (WM115), -0.00013 (NZM40), and -0.01720 (CM150-Post), respectively. Further, the 95% limits of agreement are very narrow at -0.03535 to 0.02149 (WM115), -0.05039 to 0.05013 (NZM40), and -0.05726 to 0.02287 (CM150-Post). Our Bland-Altman analysis suggests that the inter-run variation is very low for any individual data point, with <1% variation between independent technical replicates. Further, there are no identifiable trends or changes in variance with increasing average values, suggesting that there are no systematic differences between runs. In this context, the Bland-Altman analysis provides strong evidence that our sequencing results remain very consistent across independent replicates and, therefore, any observed alterations in DNA methylation level between unedited and edited samples are the result of targeted manipulation via our editing system.

### 3.4 Effective Editing Window of the dCas9-SunTag System

Though the primary objective of this study was to perform targeted manipulation of our previously identified 58 bp EBF3 locus, our DNA methylation analysis was designed to sequence a 285 bp amplicon of the EBF3 promoter, containing a total of 34 CpG sites (chr10:131,763,460–131,763,744; reference genome GRCh37/hg19). Concordant with previous findings, our dCas9-SunTag system was able to induce robust methylation change across at least our 285 bp amplicon (Figure 4); effective methylation editing has previously been described up to 1 kb from the target site using dCas9-SunTag [18]. Indeed, dCas9-SunTag systems have a broader effective editing window than other described systems [29, 40, 45–47], due to the SunTag protein scaffold tail and capacity for effector multimerisation. Due to the position of our gRNA, CpG sites 1-4 within our sequenced amplicon are located within our gRNA target sequence. We observed demethylation at these four CpG sites, regardless of using a methylating (scFv-DNMT3A) or demethylating (scFv-TET1CD) effector construct. Again, this is consistent with previous findings which suggest that binding of the dCas9 module at the target site sterically prevents the activity of effector enzymes at underlying CpGs [25]; further, bound dCas9 may preclude the action of endogenous DNMTs, thereby promoting demethylation at these CpG residues.

**Figure 4.**
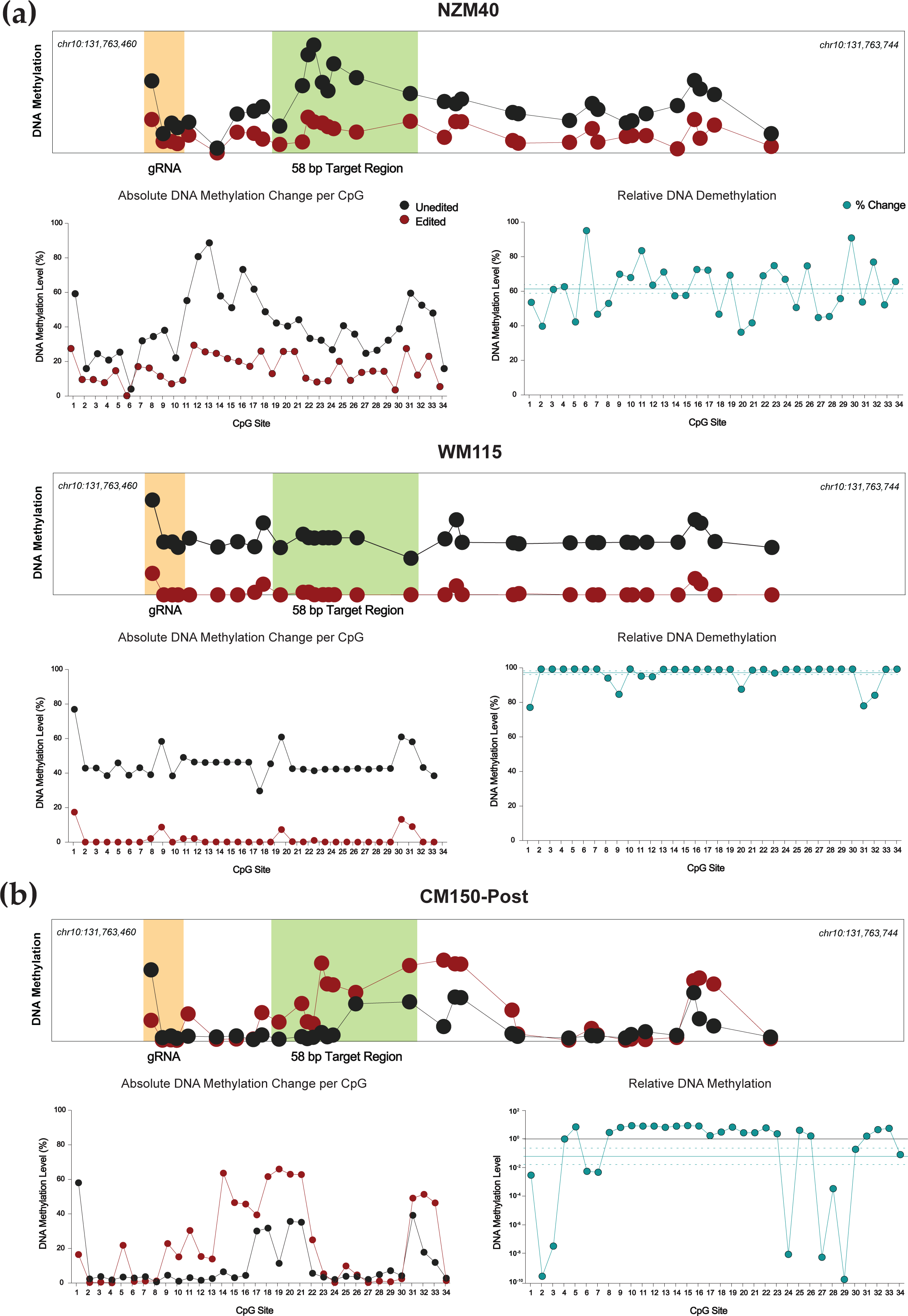
**(a)** Targeted DNA demethylation of the *EBF3* promoter region for NZM40 and WM115 cell lines. Mean DNA methylation levels between unedited (black) and edited (red) samples are plotted for each of the 34 CpG sites within the 285 bp sequenced amplicon of the *EBF3* promoter. Mean per CpG methylation levels are shown by relative genomic position (top), with the gRNA binding site (orange) and target region (light green) highlighted as shown. Absolute DNA methylation change (bottom left) and relative demethylation (%) (bottom right; teal) are plotted per CpG for the entire amplicon. Mean relative demethylation across all 34 CpGs (teal line) and standard error of the mean (SEM; dotted teal line) are shown. **(b)** Targeted methylation of the *EBF3* promoter region for cell line CM150-Post. Mean DNA methylation levels between unedited and edited samples are plotted for each of the 34 CpG sites by relative genomic position (top). Absolute DNA methylation change (bottom left) and relative demethylation (proportion) (bottom right) are plotted per CpG for the entire amplicon. Mean relative methylation across all 34 CpGs and SEM are plotted as shown.

Our dCas9-SunTag, scFv-TET1CD system demonstrated effective and consistent demethylation across the entire 285 bp amplicon (Figure 4a). For both NZM40 (mean relative demethylation 61.32%, SEM ±2.50%) and WM115 (mean relative demethylation 96.75%, SEM ±1.08%), relative levels of demethylation were remarkably consistent across all 34 CpG sites, suggesting that our editing tool has reliable efficacy at CpG residues within a locus of this size. Interestingly, dCas9-SunTag, scFv-DNMT3A was more variable, with peak methylation increases observed around 25-120 bp and 180-190 bp from the PAM site (Figure 4b). Several CpG sites within the wider amplicon displayed a decrease in DNA methylation; however, this occurred exclusively at CpG sites with comparatively low baseline methylation levels (<10%). As such, these methylation changes may carry less biological significance. The variations in editing efficacy observed with this tool may be secondary to structural differences between scFv-TET1CD and scFv-DNMT3A not permitting enzymatic activity at these residues, or due to an innate propensity for certain CpG sites to remain unmethylated. Overall, both tools were able to induce efficacious local methylation change within at least the constraints of our amplicon, with the exception of CpG sites underlying the bound dCas9 module, that underwent demethylation in a cell lines irrespective of which effector was used.

### 3.5 Assessment of On- and Off-Target Activity for Targeted Editing

Gold-standard methods for assessing off-target impacts include whole genome or ChIP-bisulfite sequencing [48, 49]. These methods, though thorough and accurate, are expensive and time-consuming. Comparatively, we employed a screening method that assesses specific loci with a high likelihood of off-target impacts, based on in silico prediction. This provided a rapid assessment of high-yield loci to give confidence in targeted editing results. As Cas9-binding at specific loci is gRNA-directed, INDEL formation at any given locus provides a surrogate measure of gRNA activity. Through selecting a smaller number of loci for assessment, this approach provides high yield off-target assessment without the need for more extensive, genome-wide approaches.

We perfomed a targeted gRNA assessment of gRNA location #1, used for editing our EBF3 target region (Figure 5). Assessment of on-target activity, as well as activity at the top ten predicted off-target loci was performed. Two negative controls (Cas9 only; gRNA only) were included. Our gRNA displayed high on-target efficacy, with 90.48% of reads in the EBF3 target region containing INDELs above background noise; comparatively, 0.46% and 0.20% of the Cas9 and gRNA negative control samples contained INDELs above background noise, respectively. All three groups had very low rates of off-target INDEL generation at predicted off-target sites, with all <1%. Given the consistency of these INDEL rates with those of both negative control samples, any off-target INDELs are likely to be due to random mutation rather than direct effects of our editing system. For targeting of the EBF3 promoter, these data demonstrate that our selected gRNA(s) are highly efficacious at the desired locus, with minimal off-target activity. These findings are consistent with previous off-target assessments of dCas9-SunTag systems, wherein whole-genome bisulfite sequencing results have demonstrated minimal off-target effects on methylation [19, 48].

**Figure 5.**
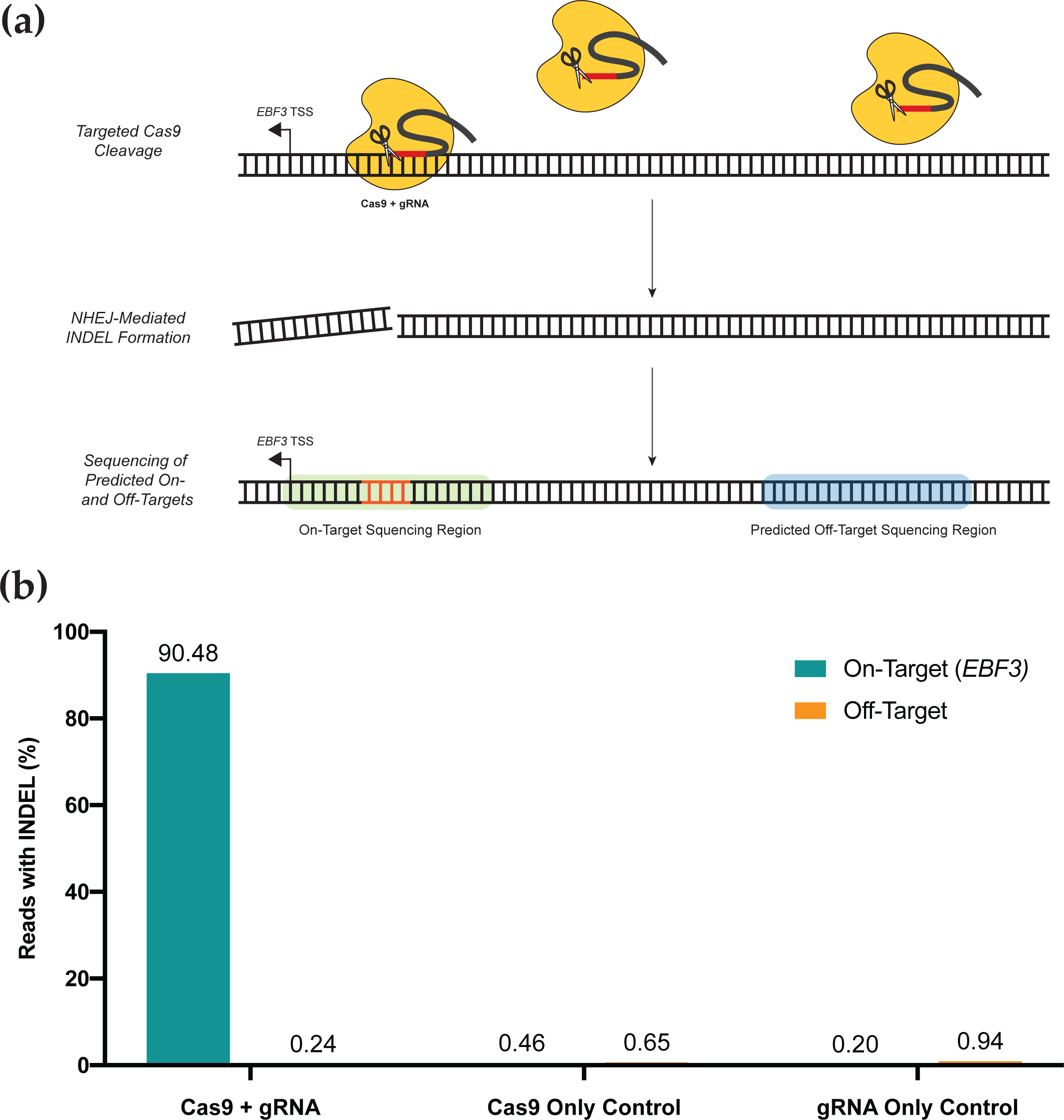
**(a)** Rapid screening method for on- and off-target assessment of unique gRNA constructs. The gRNA of interest and active Cas9 construct are transfected into cell lines. Directed by the gRNA, Cas9 binds to genomic loci and induces double-stranded cleavage of the DNA molecule, which is subsequently repaired via non-homologous end joining (NHEJ). NHEJ introduces insertions or deletions (INDELs; orange) at the repaired locus which can be detected via sequencing, providing a surrogate for gRNA + Cas9 activity at any given locus. The on-target locus and top ten off-target loci are then sequenced via targeted high-throughput sequencing. **(b)** Assessment of on- and off-target activity for the gRNA used to successfully edit *EBF3* promoter methylation. Percentage of sequenced reads containing INDELs provides the surrogate measure of activity at each locus; data from off-target loci are combined as shown. Our gRNA shows very high on-target efficacy of INDEL generation and minimal off-target impacts for the top ten predicted off-target loci. Cas9 only and gRNA only controls display minimal on- or off-target effects.

Overall, we demonstrate a novel approach for gRNA assessment, which provides not only high-yield and rapid screening for off-target effects of our editing system, but also accurately determines the on-target efficacy for a respective gRNA in a genuine in vitro context. With respect to our dCas9-SunTag system, these results provide confidence that with appropriate gRNA selection, a high level of on-target binding can be achieved, with minimal impacts on the wider DNA methylome.

### 3.6 Design and Delivery Considerations for Editing System Selection

Several CRISPR-based systems have now been described for DNA methylation editing, each providing unique advantages and limitations (Table 1). Our system is a variation of the SunTag system, originally described by Tanenbaum *et al* (2014) [50] and first adapted by Morita *et al* (2016) [18] for targeted DNA methylation editing. dCas9-SunTag systems are a widely used multimerisation approach which have shown high efficacy for both DNA methylation and demethylation, whilst displaying superior off-target profiles than other editing systems [18, 48]. SunTag systems also provide scope for the multimerisation of complementary effector constructs, such as TET1CD and VP64 for multi-level gene activation [51, 52].

Transient methods of construct delivery, including lipofection, provide a rapid and simple method of construct delivery and has been a favoured approach for DNA methylation editing to date [18, 29, 30, 39, 40, 45–47, 53, 54]. Transient delivery methods have the benefit of simplicity and cost-effectiveness, allowing for straightforward, proof-of-concept experiments to be performed in a timely manner. Indeed, stable transfection methods, especially those which rely on viral delivery, require more specialised facilities and training to be performed, as well as typically longer time frames for the generation of transduced cell lines. As such, we prefer the use of a transient delivery approach in the first instance, supplemented with more stable approaches secondarily. The major advantage of stable delivery options, interestingly, would appear to be in efficiency of delivery for difficult cell lines rather than the stability of editing. In fact, numerous studies have now demonstrated that transient delivery of DNA methylation editing systems results in stable changes in methylation for long periods, with persistent changes observed for up to 50 days in replicating cells [29, 30, 46]. There is, however, some evidence that viral delivery may allow for more substantial targeted methylation changes in the context of dCas9-SunTag, scFv-DNMT3A [19].

To enhance the efficacy of our CRISPR-based tool in melanoma cell lines, we are currently optimising a lentiviral delivery system. Lentivirus-mediated transduction of our CRISPR-based editing system has the potential to complement our current approach via increasing the efficiency of targeted methylation editing and enabling the generation of stably expressing cell lines for further applications.

Our own data support dCas9-SunTag as an effective system for manipulating DNA methylation. For the first time in human melanoma cell lines, we demonstrate highly efficacious, site-specific DNA methylation editing of the EBF3 gene promoter; we also demonstrate both targeted methylation and demethylation of a single locus. Paired with scFv-DNMT3A, our dCas9-SunTag-based editing system was able to induce a substantial relative increase in DNA methylation at our target locus, in a lowly-methylated cell line. Likewise, our scFv-TET1CD effector was able to achieve up to near-complete demethylation, or unmethylation, in moderate to highly-methylated cell lines. Impacts on DNA methylation were consistent across our entire sequenced amplicon, providing support for the relatively broad editing window of this tool. Further, these effects were rapidly achieved with transient delivery our editing system. In addition to high levels of on-target efficacy, our off-target screening demonstrates that these tools act with high specificity, displaying minimal off-target activity. Together, these data highlight the efficacy, specificity and flexibility of our described system, which can be adapted for both targeted DNA methylation or demethylation using this fast and straighforward approach.

### 3.7 Limitations and Troubleshooting

Though optimized to suit the human melanoma cell lines WM115, CM150-Post and NZM40, this protocol may also be broadly applicable to other human cell lines for which Lipofectamine 3000 is an effective transfection reagent, particularly human melanoma cell lines. In our experience, electroporation proved an ineffective method for delivery of our editing system in these particular cell lines (data not shown) and so lipofection was chosen as our preferred method for transient construct delivery.

#### 3.7.1 Timing of Transfection

The timing of cell culture in relation to transfection is crucial to maintain cell health, ensure adequate cell numbers are available, and to maximise transfectability. Different cell lines may require different growth times to achieve this, and so, a trial of growth from frozen stocks to adequately cultured cells is recommended prior to attempting transfection.

#### 3.7.2 DNA Quality and Total DNA Input

Before beginning the first transfection, or when generating new batches of plasmid DNA for transfection, we recommend screening the isolated DNA for accuracy. As each of the constructs is produced using restriction cloning, this can easily be done via a restriction digest (e.g., RsrII/SpeI digest of an effector construct). Additionally, transfection is better tolerated when DNA purity is high and so a DNA purity assessment may be valuable prior to transfection (i.e., ensuring an adequate A260/A280 ratio).

Though optimized for the cell lines WM115, CM150-Post and NZM40 to 1500 ng total DNA per transfection reaction, optimal DNA input may vary between cell lines. If excessive cell toxicity is observed using 1500 ng of total DNA, we recommend optimizing DNA input for each respective cell line.

#### 3.7.3 Exposure to Transfection Reagents

Similar to total DNA input, Lipofectamine 3000 concentration and exposure time to transfection reagents are key factors which we have optimized for cell lines WM115, CM150-Post and NZM40. Therefore, if cell toxicity is high, or if using different cell lines, we recommend optimizing Lipofectamine 3000 concentration per reaction and/or exposure time to transfection reagents. Each of these optimization experiments can be performed via the described transfection method and FACS selection, with subsequent analysis of the FACS data output to determine optimal transfection conditions. Note, a reverse-transfection method is described here as our group has had previous success using reverse transfection for these adherent cell lines. It would be reasonable to trial a standard transfection method in other cell lines, which has been demonstrated previously [13].

#### 3.7.4 Spectral Overlap Considerations for FACS

If spectral overlap is predicted, then we recommend performing a FACS compensation experiment using cells transfected with each respective plasmid, individually. This allows the FACS system to be adjusted for inherent overlap between the emission spectra of fluorophores used in this editing system. To differentiate live versus dead or dying cells in the APC-Cy7 channel, we recommend using LIVE/DEAD Fixable Near-IR Dead Cell Stain, as per the manufacturer’s instructions. If the spread of live versus dead cell populations shows excessive overlap, however, we recommend using a more dilute stain preparation (i.e., 1 µL stain per 6-8 mL 1 x DPBS). This may need to be modified with different cell lines.

#### 3.7.5 Transfection Efficiency

Known limitations of our approach include relatively low transfection efficiency and, subsequently, low cell output post-FACS, largely due to the difficulties of co-transfecting three large plasmids simultaneously. Low cell outputs may present challenges for downstream applications, including investigation of changes in gene expression or chromatin organisation. For cell lines where cell numbers are insufficient for downstream analysis, alternative approaches should be considered, including other methods of transient system delivery. Without employing a stable delivery method such as viral packaging, substantial improvements in transfection efficiency may be difficult to achieve for some cell lines. Furthermore, the use of combined, or smaller, constructs could potentially mitigate the difficulties of co-transfecting multiple large plasmids [18]. When considering further analysis of samples with low cell numbers, we recommend the use of specialized kits for low cell input (e.g., EZ DNA Methylation-Direct Kit).

## 4. Conclusions

Here, we describe an effective protocol for targeted DNA methylation editing in human melanoma cell lines. We use transient lipofection as the delivery strategy for our dCas9-SunTag-based editing system and provide a thorough workflow, including: system design; delivery; analysis; and a novel method for rapid off-target assessment. Although we focus on editing DNA methylation in the mammalian genome, the steps and principle of this protocol could be used to edit and study methylation of other model organisms, such as zebrafish [55, 56]. Employing this dCas9-SunTag-based editing approach, we performed highly efficacious manipulation of DNA methylation within the *EBF3* promoter, across multiple cell lines, inducing substantial levels of targeted methylation and demethylation, with minimal off-target effects.

## 5. Data Availability Statement

The original contributions presented in the work are included in the article. Further inquiries can be directed to the corresponding authors.

## 6. Author Contributions

Conceptualization, A.C, R.J.W; methodology, G.G, A.C, R.J.W; software, J.S, R.B, R.W, A.U, G.G and R.J.W; validation, J.S, R.B and R.W; formal analysis, J.S, R.B, R.W, A.U, G.G, R.D and R.J.W; investigation, J.S, R.B, R.W, A.U, R.D, R.J.W; resources, R.D, M.R.E, A.C and R.J.W; data curation, J.S, R.B, R.W, A.U, G.G and R.J.W; writing—original draft preparation, J.S; writing—review and editing, all authors; visualization, J.S, R.B, R.W, A.U, G.G and R.J.W; supervision, A.C, R.J.W and M.R.E; project administration, A.C and R.J.W; funding acquisition, A.C. All authors have read and agreed to the published version of the manuscript.

## 9. Acknowledgments

We would like to acknowledge Michelle Wilson (Flow Cytometry Technician, University of Otago) for her expertise and assistance in performing FACS, Dr Jisha Antony for her helpful discussion of CRISPR experiments, and Professor Ian Morison and Dr Peter Stockwell for their support in our work on methylation editing.

## 11. Appendix A

Reagents:

- Human melanoma cell lines to be transfected (e.g. NZM40, WM115, CM150-Post)
- Appropriate cell culture medium for chosen cell line(s)
- 0.05% Trypsin-EDTA
- BsmBI restriction endonuclease and NEBuffer 3.1™ (New England Biolabs)
- Shrimp alkaline phosphatase (rSAP)
- DNA Clean and Concentrator-5 Kit (Zymo Research), or equivalent
- T4 DNA Ligase and 10x T4 DNA Ligase Buffer (ThermoFisher Scientific)
- Pre-prepared CRISPR-methylation plasmids
- LB agar and broth for propagation of pre-prepared CRISPR-methylation plasmids
- Appropriate antibiotic for plasmid selection (e.g. 100 µg/mL ampicillin)
- Plasmid isolation system (e.g. GenCatch™ Plasmid Plus DNA Maxiprep Kit (Epoch Life Science Inc))
- Opti-MEM serum-free medium (Invitrogen)
- Lipofectamine 3000 reagent and P3000 reagent (Invitrogen)
- 1 x Dulbecco’s phosphate buffered saline (DPBS)
- Sterile autoMACS buffer (1 x DPBS + 1% FCS + 2 mM EDTA)
- Ice for keeping cells pre- and post-fluorescence-activated cell sorting (FACS)
- DNA isolation kit for transfected FACS-selected cells
- Bisulfite conversion kit (e.g. EZ DNA Methylation-Gold Kit (Zymo Research)
- For low cell numbers, the combined DNA isolation and bisulfite conversion kit (e.g. EZ DNA Methylation-Direct Kit (Zymo Research))
- Reagents for targeted DNA methylation analysis (see 2.5 Targeted DNA Methylation Analysis for Confirmation of CRISPR-Editing)

Equipment:

- Access to the Benchling online platform for gRNA Design (http://benchling.com)
- Cell culture facilities and incubator with appropriate conditions for chosen cell line(s)
- Cell culture flasks
- Tube(s) for cell resuspension (e.g. 15 mL or 50 mL Falcon conical centrifuge tubes)
- Centrifuge suitable for 15 mL or 50 mL Falcon tubes
- Cell counting apparatus (e.g. haemocytometer)
- Shaking incubator suitable for mid-scale (200-500 mL) bacterial cultures
- 1.5 mL microcentrifuge tubes
- 6-well cell culture plates
- 15 mL Falcon tubes for FACS cell preparation and collection
- FACS system with capacity for sorting cells which are positive for at least three fluorophores simultaneously (e.g. BD FACSAria Fusion (BD Biosciences))
- Equipment for targeted DNA methylation analysis (see 2.5 Targeted DNA Methylation Analysis for Confirmation of CRISPR-Editing)
- Access to the NCBI Primer-BLAST online tool for optional gRNA assessment experiments (https://www.ncbi.nlm.nih.gov/tools/primer-blast/index.cgi)

## 12.

**Supplementary Table S1.**
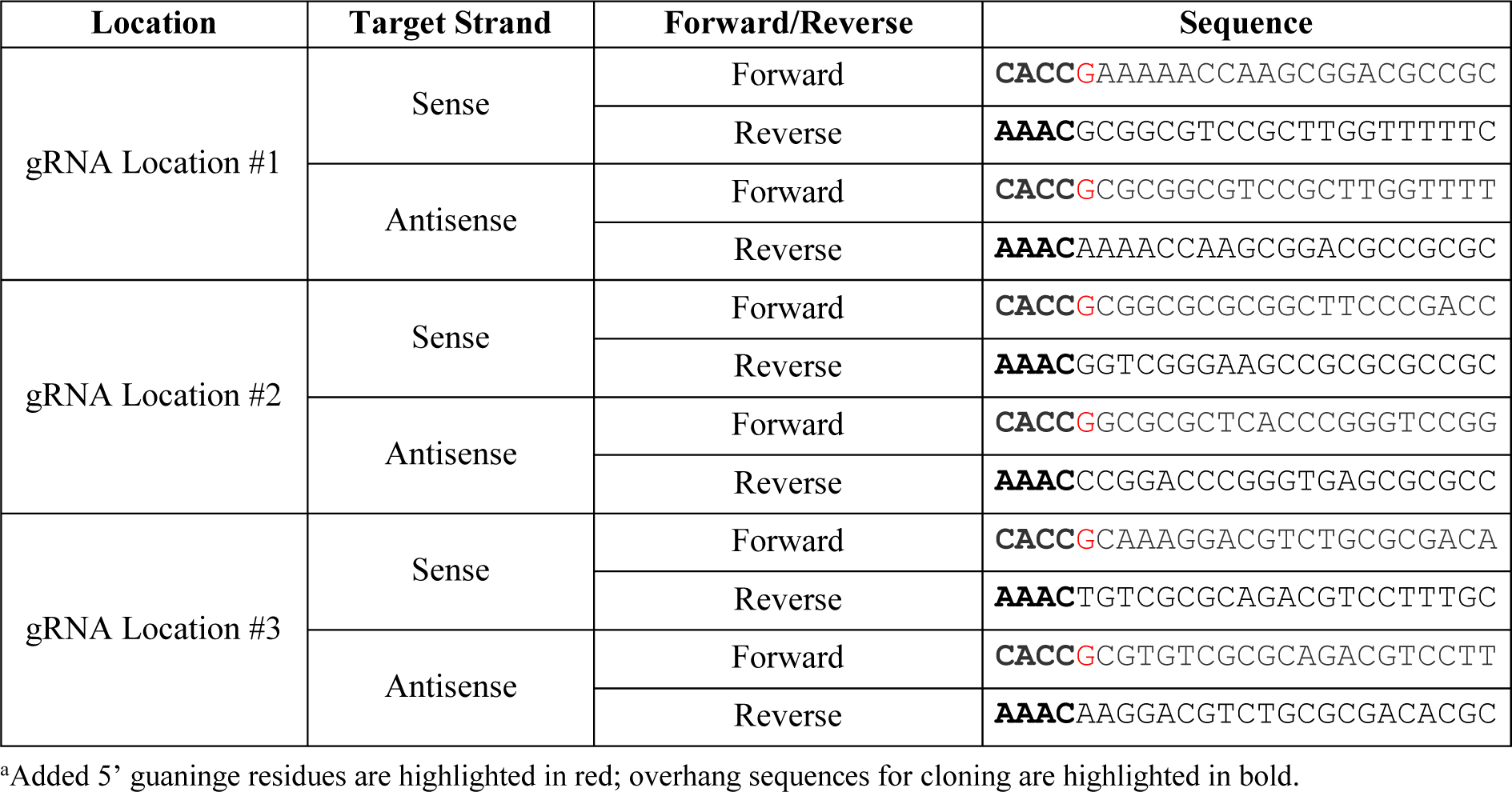
Forward and reverse oligonucleotide sequences for *EBF3* promoter gRNAs.

## 13. Figure Legends

**Supplementary Figure S1.**
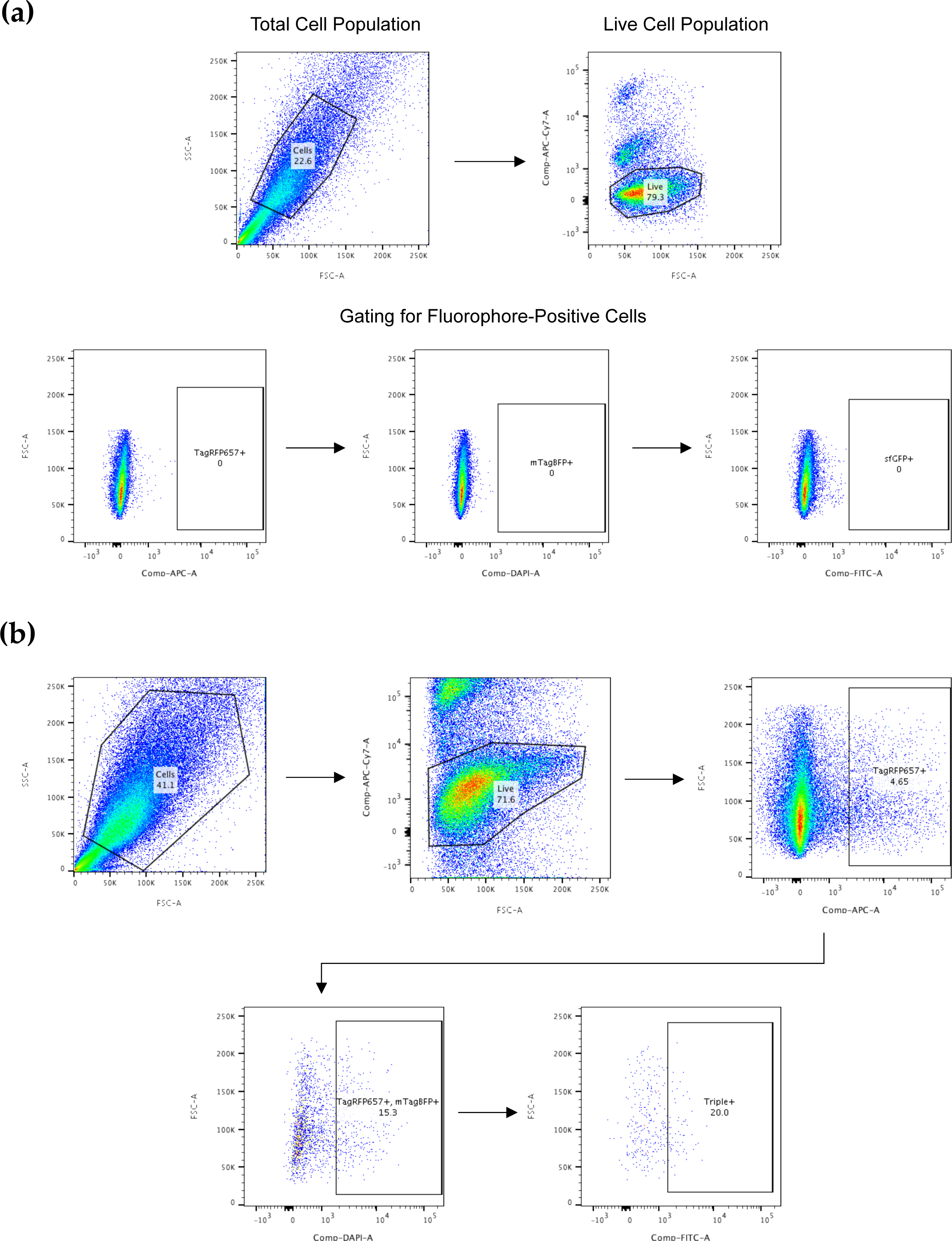
FACS for CRISPR-edited cells. **(a)** Gating setup using DNA-free negative control samples. Shown is the use of a DNA-free negative control sample to set accurate gates for fluorophore-positive (i.e., transfected) cells. Gates are set for each respective channel to ensure that only genuine fluorophore-positive cells will be sorted during FACS. **(b)** Example of FACS for CRISPR-edited NZM40 human melanoma cells. Shown is an example of the sequential gating strategy for sorting triple-positive transfected cells in practice, which selects sequentially for: the true cell population; live cells only; cells expressing TagRFP657; cells also expressing mTagBFP; cells also expressing sfGFP. Hence, all cells which are eventually sorted into the final collected population are live, triple-positive cells.

**Supplementary Figure S2.**
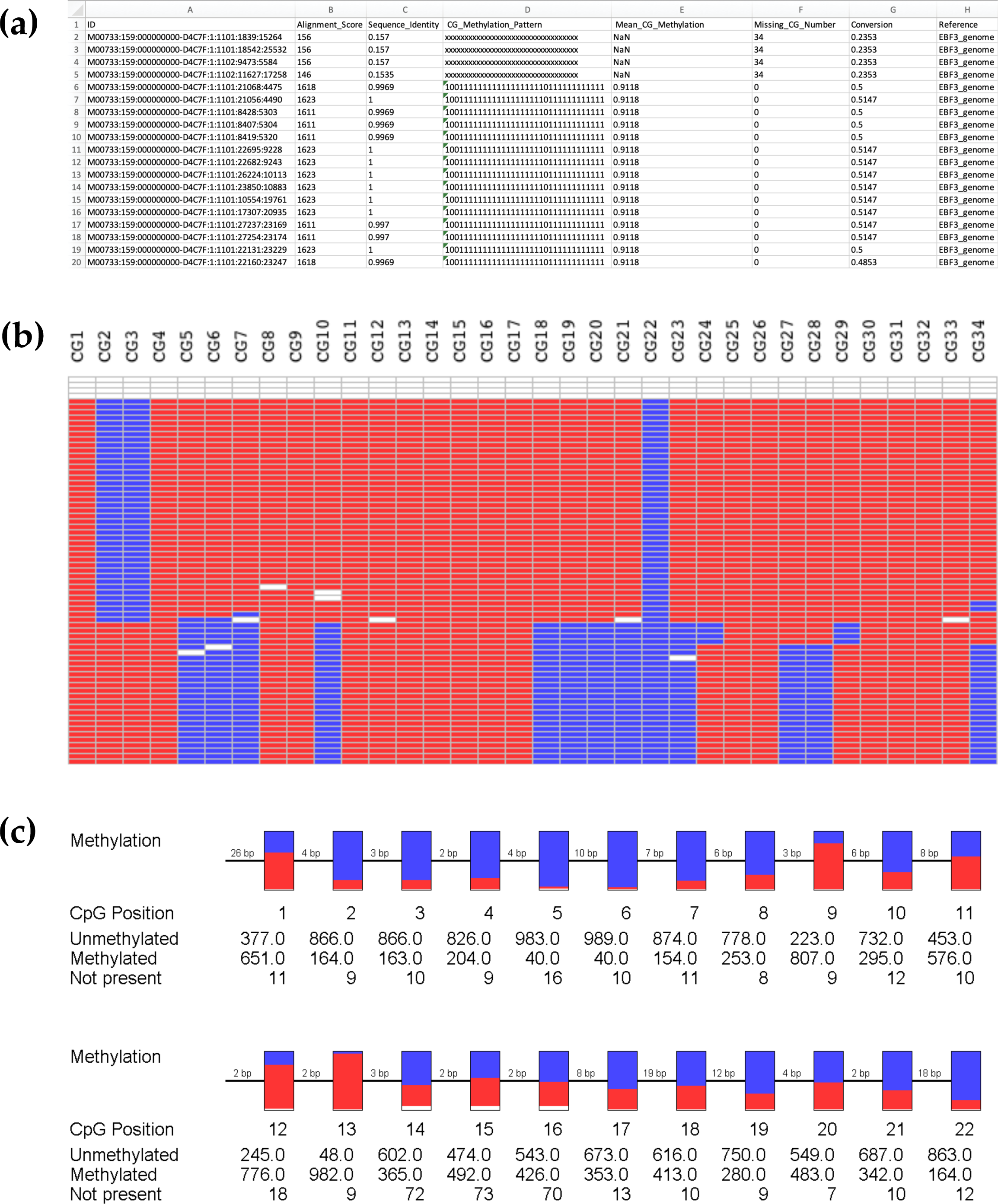
BiQ Analyser HT processing during the analysis of targeted methylation sequencing data. **(a)** Example output results.tsv file (opened in Microsoft Excel) summarising the results of sequence alignment for a particular demultiplexed sample with 34 CpG sites interrogated in BiQ Analyser HT. This provides all information for a particular read following alignment and calculates multiple parameters to assess the success of read alignment. The binary methylation output is also shown (1 = methylated; 0 = unmethylated; x = unaligned). As shown, the first four reads in this example have very low alignment scores, and these will be removed by further analyses (alignment score threshold ≥1000). **(b)** Example methylation heatmap output visually displaying binary methylation data for 34 CpG sites and for each sequenced read. Each row represents an individual sequencing read, whilst each column reflects a different CpG within the sequence as labelled. Methylated CpG sites are labelled red; unmethylated sites in blue; and unaligned in white. **(c)** Pearl necklace plots, again visualising example methylation data per CpG site. However, rather than separated reads, these plots illustrate the proportionate methylation level at each CpG, across all sequenced reads for that sample, with the number of unmethylated, methylated and ‘Not present’ occurrences shown. Note that the pearl necklace plot shown in (c) does not correspond directly to the data shown in (b). This is because the heatmap shown in (c) only shows a portion of the total data set to provide an example of a methylation heatmap and (c) only shows an example of the first 22 CpG sites.

